# Integrated *ex vivo* screening and transcriptomic profiling to prioritize drug combinations for rare cancers

**DOI:** 10.64898/2026.07.28.741264

**Authors:** Ritu C. Shah, Alex T. Larsson, Belinda B. Garana, Yang Lyu, Zachary Seeman, Elizabeth Rono, Kyle B. Williams, Dana Borcherding, Kangwen Xiao, Xiaochun Zhang, Yannick Mahlich, Jeremy Jacobson, Lisa B. Fridman, Yan Zhou, Christine A. Pratilas, Angela C. Hirbe, David K. Wood, David A. Largaespada, Sara J.C. Gosline

**Affiliations:** Masonic Cancer Center and Department of Biomedical Engineering, University of Minnesota, MN, 55455; Masonic Cancer Center and Department of Pediatrics, University of Minnesota, Minneapolis, MN, 55455; Biological Sciences Division, Pacific Northwest National Laboratory, Seattle, WA 98109; Division of Oncology, Washington University in St. Louis, St. Louis, MO; Department of Oncology, Sidney Kimmel Comprehensive Cancer Center at Johns Hopkins, Baltimore, MD; Department of Laboratory Medicine and Pathology, University of Minnesota, MN, 55455; Department of Biomedical Engineering, Oregon Health and Sciences University, Portland, OR

**Keywords:** 3-dimensional (3D), malignant peripheral nerve sheath tumor (MPSNT), drug screen, *ex vivo*, patient derived xenograft (PDX)

## Abstract

Discovering effective drug combinations requires testing many dose combinations across a diverse panel of tumor models. This approach is limited in rare cancers by a scarcity of cell lines and representative high-throughput models that would make exhaustive screening feasible and predictive. Patient-derived xenograft (PDX) models are genomically representative but too low-throughput for this purpose. Culturing PDX cells *ex vivo* in three-dimensional (3D) matrices offers a genomically representative and clinically relevant platform for preclinical drug testing, capturing the microenvironmental cues that shape *in vivo* drug response while requiring only limited tissue per assay. Toward this end, we designed and validated an experimental-computational framework, “*ex vivo* assessment of combination therapies” (EXACT), to enable drug combination discovery in rare tumors. Using PDX models of malignant peripheral nerve sheath tumors (MPNST), we built a platform to culture PDX cells *ex vivo* over multiple days, monitoring drug sensitivity and measuring transcriptomic responses to treatment. Computational analysis of these transcriptomic responses then identifies which compensatory pathway creates a unique vulnerability to a second drug. EXACT thus offers a biologically informed, scalable approach for prioritizing drug combinations in rare tumors, nominating drugs alongside biological rationale. Using this methodology, we identified a MEK inhibitor plus HDAC inhibitor combination with enhanced activity *in vitro* and *in vivo*, forming the basis of an active clinical trial. This platform could be adapted for real-time use with primary patient specimens, enabling personalized therapeutic discovery.

**Significance:** EXACT integrates PDX-derived 3D drug screening with biologically informed computational analysis to identify and explain effective combinations, providing a scalable strategy for therapeutic discovery in rare cancers such as MPNST.

## Introduction

Despite the clinical success of many precision targeted cancer therapies, treatment-emergent drug resistance and metastasis remain major obstacles to durable treatment responses and cures^1^. This challenge motivates interest in combination therapies, but finding drugs that are effective in combination requires understanding both the biological and functional impact of each drug individually to identify pairs of drugs that can be more efficacious together^2^. Typically, established cancer cell lines grown in two-dimensional (2D) cultures on plastic are the workhorse for this work, enabling genomic and functional profiling across diverse cancer types and the construction of publicly available drug response datasets ^2–6^. Computational tools built on these datasets have been developed to predict effective drug combinations from untreated samples^7^. However, these models fail to generalize to rare tumors, where established cell lines are often lacking because they do not predict drug response well enough.

Malignant peripheral nerve sheath tumors (MPNST), aggressive sarcomas that arise commonly in individuals with neurofibromatosis type 1 (NF1) ^8,9^, exemplify this challenge. Current treatment options are limited, with no FDA-approved targeted therapies to date ^10,11^ despite clinical trials that attempted to translate laboratory based discoveries. Five-year survival for metastatic or recurrent disease remains below 20%, underscoring the urgent need for new treatment strategies guided by molecular insights ^9,12–14^. As with other rare cancers, established cell line models may fail to capture the genomic heterogeneity and don’t allow for analysis of stromal interactions, and drug screening efforts have yet to yield clinical benefit.

For rare cancers such as MPNST, the limited number and molecular diversity of available cell lines constrain their ability to represent the broader patient population. Patient-derived xenografts (PDXs) help address this limitation by maintaining key genomic and epigenomic features of the original tumors^15,16^, but are disadvantaged by their low-throughput nature, time intensive requirement, and use of large numbers of animals to achieve translationally relevant datasets. Several *ex vivo* culture systems, including organoids, microtissues, and hydrogels, have been shown to preserve primary patient tissue and the genomic landscape of the source material, making PDX-derived tumor cells an attractive source for drug testing ^17–19^. We recently developed a matched PDX and *ex vivo* culture platform to identify molecular biomarkers of drug response and enable personalized therapies ^20^. However, even with current PDX resources, limited tumor yield per model and the material demands of *in vitro* assays make exhaustive screening of drug doses and combinations across multiple patient-derived models impractical. These experimental constraints highlight the need for computational approaches that can identify and prioritize the most promising drug combinations for experimental validation.

Computational models that integrate omics data with drug sensitivity measurements have been developed to address this prioritization challenge, with large cell line datasets such as CCLE, GDSC, and DepMap providing the foundation for biomarker discovery and drug response prediction^3–6^. Machine learning and network-based approaches trained on these data have shown promise in stratifying patients, nominating drug targets, and uncovering mechanisms of resistance (e.g. Kuenzi et al) ^2^. Such approaches include network propagation, matrix factorization, and deep learning-based models^21^. However, most efforts emphasize single-agent activity, and tools that do predict combinations generally focus on drug synergy, which is uncommon and requires large training datasets ^4,7,22^. Even the best-performing combination prediction models failed to identify 20% of effective combinations in a global benchmark analysis ^23^. In the rare tumor setting, where false negatives carry high cost, existing frameworks are insufficient. What is needed instead are approaches that can simultaneously model drug activity, capture network adaptation to drug exposure, and prioritize combinations for experimental testing.

To address this challenge, we developed EXACT (*ex vivo* assessment of combination therapies), an integrated experimental–computational framework that leverages PDX models and the 3D-Matrix Embedded Drug Screening (3D-MEDS) *ex vivo* culture platform to nominate and validate effective drug combinations in a physiologically relevant context (**Fig. 1**). Building on our previous work, EXACT simultaneously captures baseline gene expression to model single-agent drug activity, measures transcriptional responses to drug treatment to characterize network adaptation, and uses both to prioritize combinations for experimental validation^20^. We demonstrate the utility of this approach in MPNST, identifying enhanced efficacy of MEK and HDAC inhibitor combination therapy, which forms the basis for an actively recruiting clinical trial (NCT06693284) developed by our team. Together, these results establish a generalizable framework for integrating computational prediction with 3D *ex vivo* drug screening to accelerate discovery of rational combination therapies for aggressive and understudied cancers.

**Figure 1.**
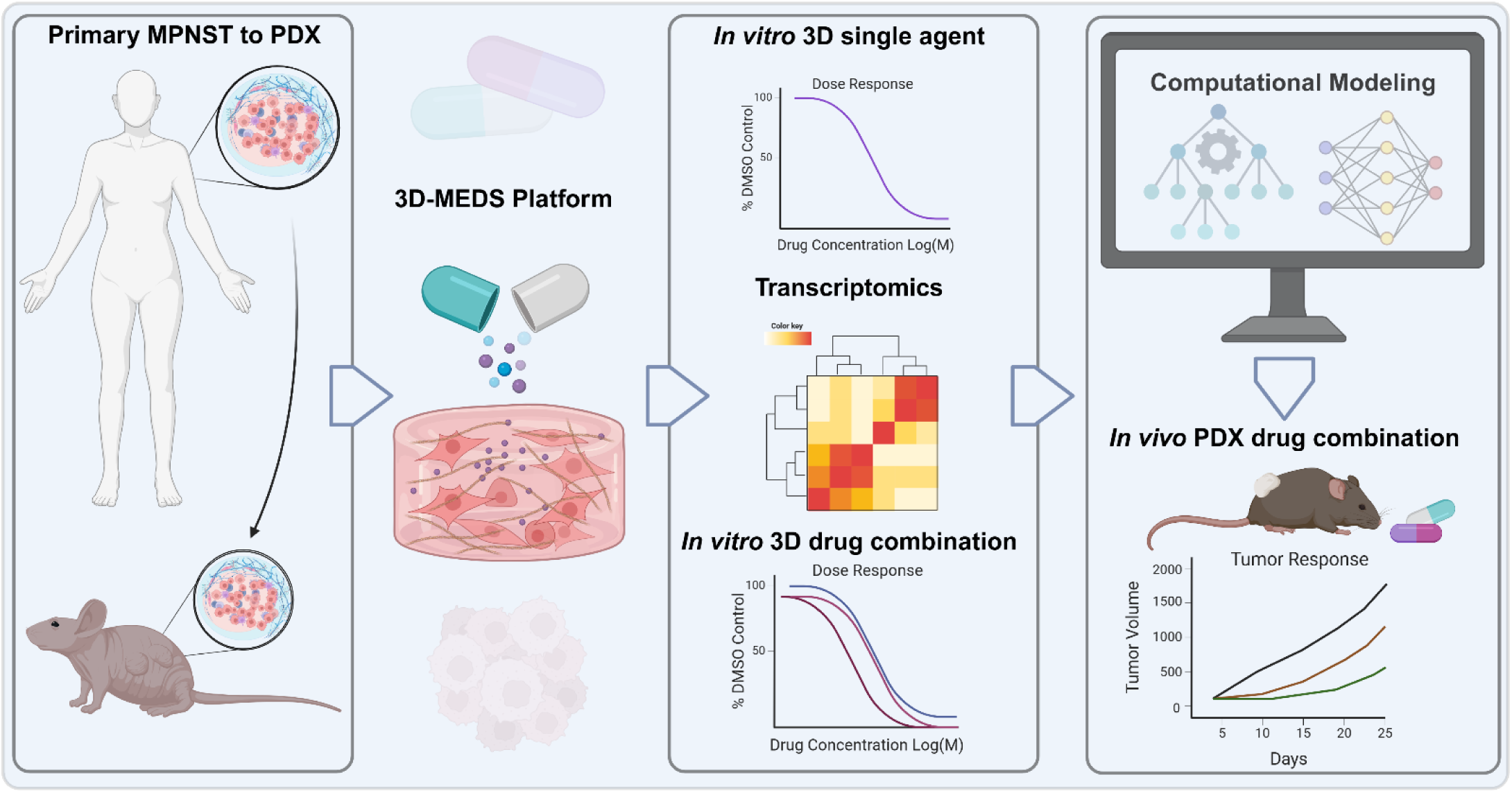
EXACT platform enables high-throughput drug screening. Overview of the *ex vivo* assessment of combination therapies (EXACT) platform. Patient derived xenografts removed from mice are plated in an ECM-rich environment creating a three-dimensional matrix embedded drug screening (3D-MEDS) platform. Cellular responses to single drug-treated samples are used to predict combination therapies via computational modeling. Combination therapies are then tested in the 3D-MEDS platform and validated *in vivo*.

## Results

### 3D-MEDS enables high-throughput drug screening in PDX-derived tumor cells

The 3D-Matrix Embedded Drug Screening (3D-MEDS) platform was designed for dissociated MPNST PDX cells using a 96-well format compatible with standard laboratory equipment and requiring minimal manual handling. Rather than microfluidic droplet generation, we used bulk basement membrane matrix encapsulation to allow simultaneous preparation of uniform 3D cultures without specialized equipment (see Methods) ^20^. This bulk-gel approach improves reproducibility and reduces shear stress relative to suspension or droplet cultures, and cell viability is quantified using luminescence-based assays that eliminate the imaging and segmentation steps required in microfluidic systems. To validate this platform, we assembled six MPNST PDX in 3D-MEDS and found that all six maintained viability through day five, with most showing increased luminescence at day two or day five, consistent with active proliferation (**Fig. S1**).

### Single-agent screening generates a functional drug response landscape across MPNST PDX

Using this platform, we tested 18 compounds from 13 drug classes across six MPNST PDX (**Table 1**), including a newly established model, MN-4, which shows retained H3K27me3 staining, suggesting intact PRC2 function (**Fig. S2**). Compounds were selected based on prior implication in MPNST signaling and known mechanisms of action involving ERK, PI3K, and epigenetic regulatory pathways ^8^. Each drug was tested across an eight-point dose-response curve using four-fold serial dilutions spanning a concentration range above the reported human maximum plasma concentration (C_max_). We quantified cell viability at each dose and summarized response using both AUC and viability at C_max_, the latter providing a clinically anchored benchmark for compounds where C_max_ values are available^24^. Both metrics were strongly correlated overall (Spearman rho=0.801, p<2.2e-16), though discrepancies proved informative in some cases. For example, palbociclib showed a low AUC suggesting anti-proliferative activity, but viability at C_max_ (**Fig. 2C, yellow vertical line**) revealed its apparent activity was driven by supratherapeutic concentrations making it among the least effective compounds. Given the strong overall agreement between metrics and the absence of published C_max_ values for some compounds, we used AUC for downstream analyses. By both measures, the HDAC inhibitor vorinostat had the greatest anti-proliferative effect across six MPNST PDX at two timepoints (**Fig. 2A, B**).

**Figure 2.**
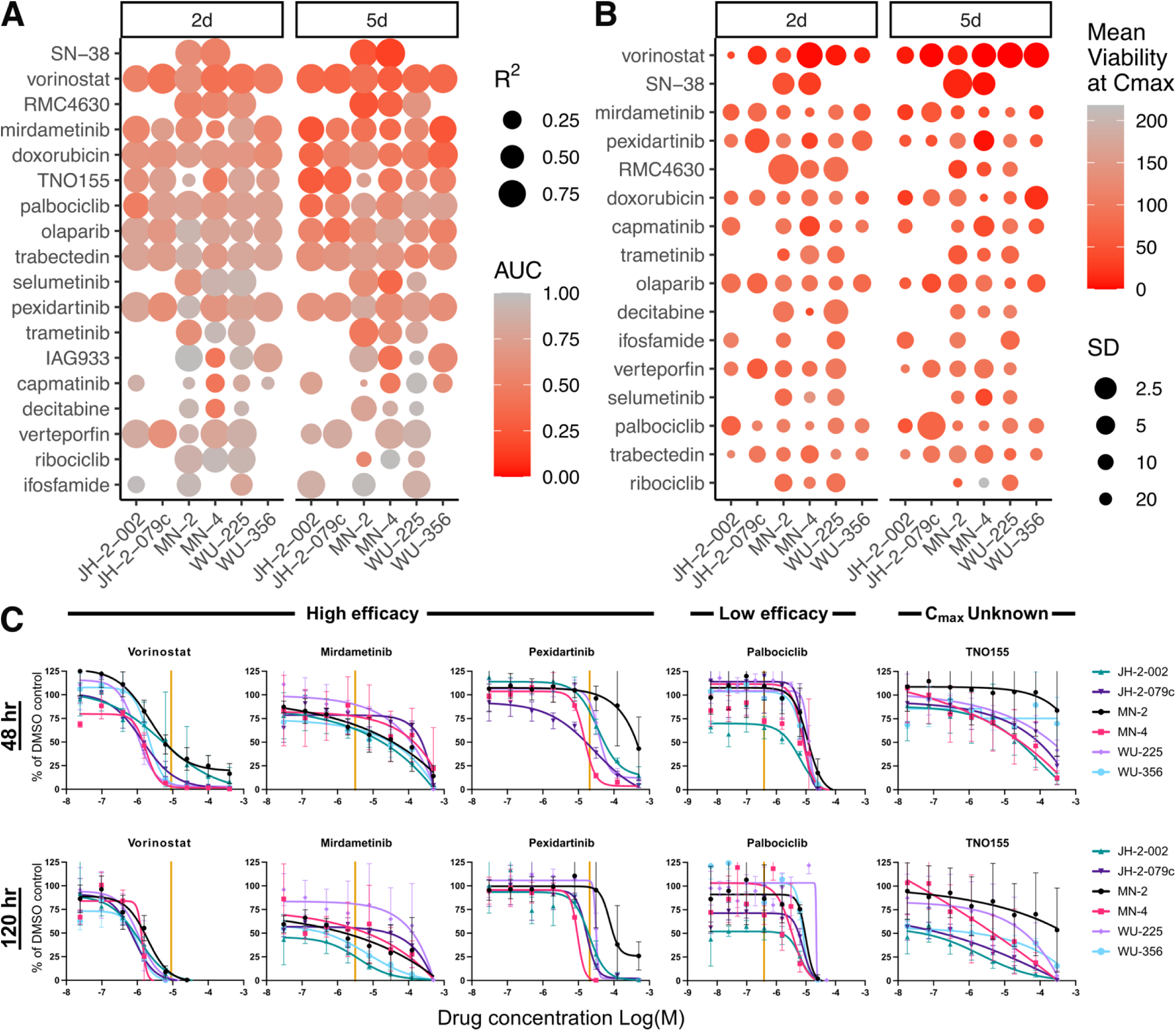
Single agent viability screening with the 3D-MEDS platform. **(A)** Area under the curve (AUC) was determined after 48 and 120 hours of individual drug treatment in six *ex vivo* MPNST PDX 3D-MEDS samples ranked by mean values. **(B)** Percent viability at biologically relevant human C_max_ doses of individual drugs was determined after 48 and 120 hours of treatment in six *ex vivo* MPNST PDX 3D-MEDS samples ranked by mean values. Size: inverse square root of standard deviation (SD). **(C)** Dose response curves of select high and low efficacious drugs in six *ex vivo* PDX 3D-MEDS show vorinostat, pexidartinib and mirdametinib perform well while palbociclib shows low efficacy as a single agent at each drug’s respective C_max_ concentration. Cell viability was calculated by normalizing luminescence values to vehicle-only control after subtracting background. Data points represent mean of triplicate values ± SD. Yellow vertical line represents C_max_

**Table 1.**
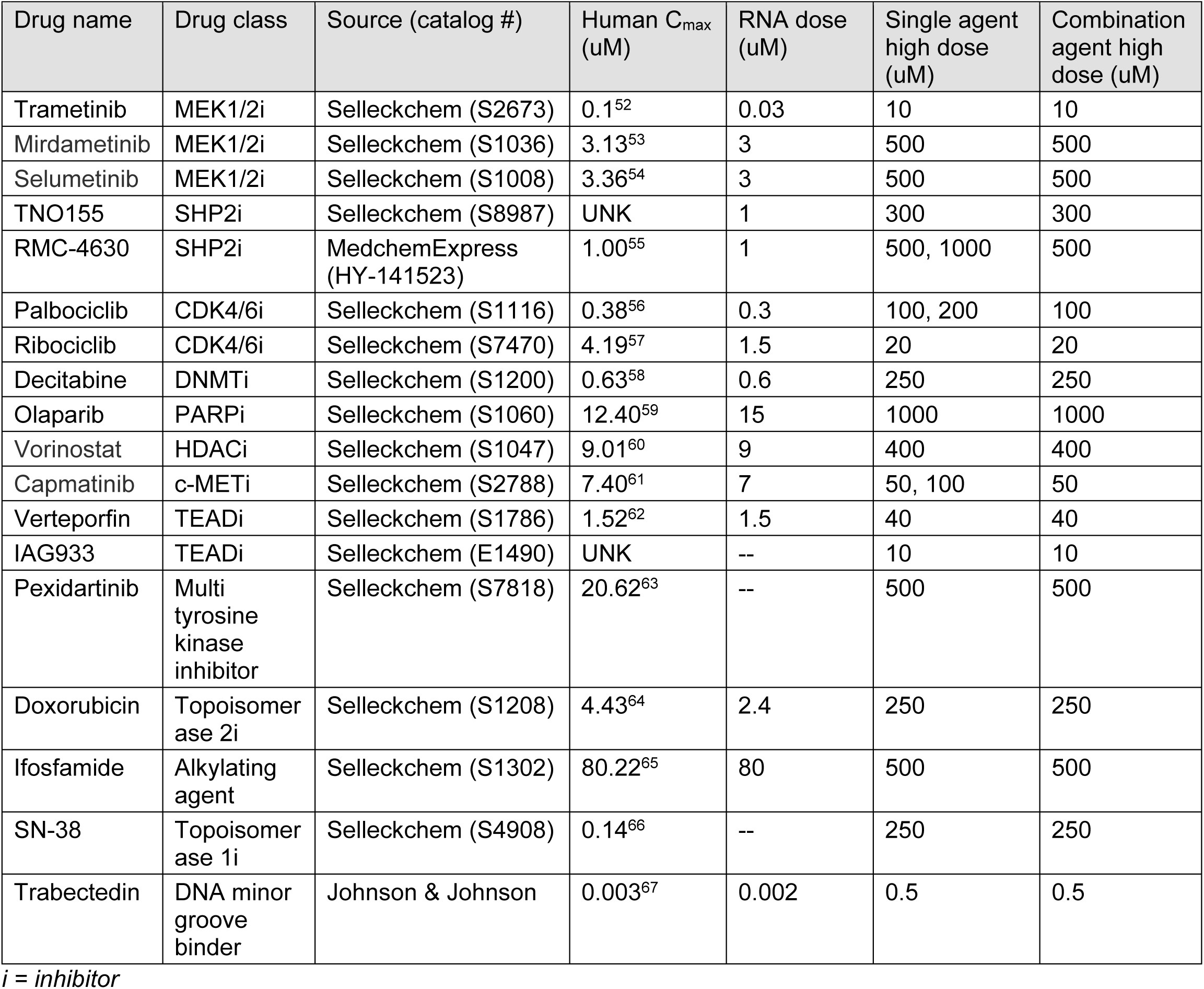
Drug information list.

### Baseline and drug-induced gene expression identify pathways linked to drug sensitivity

To ask whether baseline gene expression could predict drug sensitivity across the molecular heterogeneity of our six MPNST PDX, we applied single-sample Gene Set Enrichment Analysis (ssGSEA) to score each untreated PDX across the 50 Cancer Hallmark pathways^25^. Spearman rank correlation between pathway scores and single-agent AUC values identified pathways negatively correlated with drug response, indicating that higher baseline expression of these pathways is associated with greater drug sensitivity (**Fig. 3A**).

**Figure 3.**
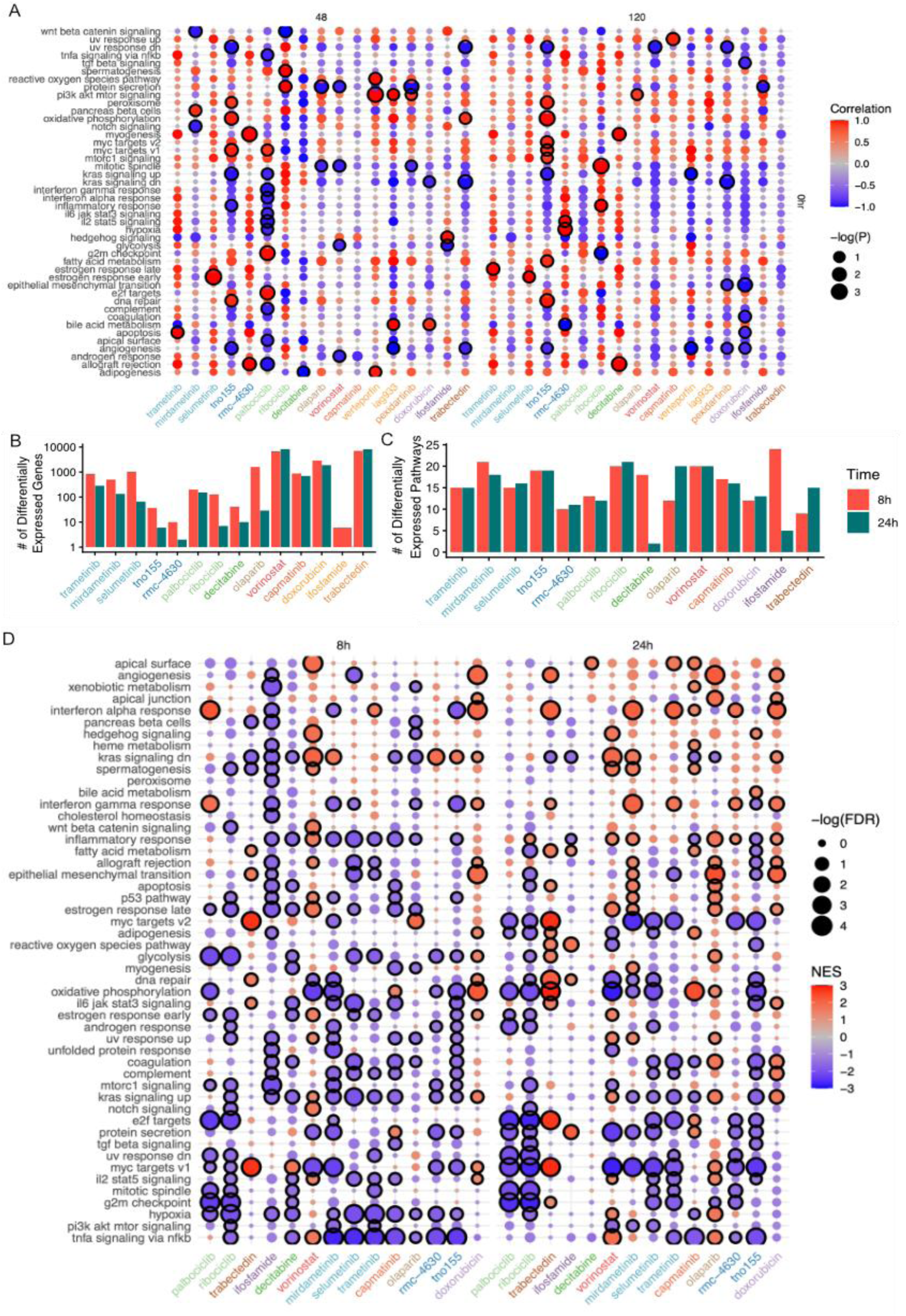
Integration of gene expression with drug sensitivity identifies potential drug combinations. **(A**) Pearson correlation between pathway scores for untreated MPNST PDX (y-axis) and the AUC values from single agent drug treatment (x-axis) measured at 48 hour (left) and 120 hours (right). Those correlations where p<0.05 are surrounded by black circle. **(B)** Number of differentially expressed genes in three MPNSTs after 8 (pink) or 24 (turquoise) hours of treatment of each of the 14 drugs in this study, colored by drug class (Table 1) **(C)** Number of differentially enriched pathways after eight (pink) or 24 (turquoise) hours of drug treatment **(D)** Statistical enrichment of top five Cancer Hallmark pathways for each drug treatment (ranked by absolute NES for each drug treatment) across different drug treatments (x-axis) and pathways and treatment time (labeled across top). Enrichment score indicated by color while significance is indicated by size. Pathways with an FDR<0.25 are outlined.

We then measured transcriptional responses to drug treatment to capture network adaptation upon single-agent exposure. Gene expression was profiled across three MPNST PDX derived cell groups (MN-2, WU-225, and JH-2-002) by RNA sequencing at 8 and 24 hours following treatment at doses near the respective human C_max_, compared to DMSO controls. Drug-induced transcriptional changes were broad but variable across compounds. Trabectedin (DNA minor groove binder) and vorinostat treatments resulted in the greatest number of differentially expressed genes, while TNO155 (SHP2 inhibitor) and ifosfamide (DNA alkylating agent) elicited the lowest number (**Fig. 3B**, adjusted p-value < 0.05; **Table S2**).

We then used ssGSEA to identify specific biological pathways altered by each drug at each time point. At the pathway level, we observed differential enrichment across most (49 out of 50) of the Cancer Hallmark pathways across all drug treatments and timepoints (p-value < 0.05 & FDR < 0.25; **Table S3**, **Fig. 3C-D**) ^26^. The most commonly upregulated pathway across the three samples was epithelial-mesenchymal transition (EMT), which is upregulated by palbociclib and ribociclib (CDK4/6 inhibitors), vorinostat, mirdametinib (MEK inhibitor), olaparib (PARP inhibitor), RMC-4630 (SHP2 inhibitor), and doxorubicin (topoisomerase 2 inhibitor). This same pathway is also significantly downregulated by ifosfamide (Alkylating agent), decitabine (DNMT inhibitor), and capmatinib (MET/ multi-tyrosine kinase inhibitor). EMT pathway activation has been shown to occur in response to many of these drug treatments, including CDK4/6, HDAC, MEK, PARP, and SHP2 inhibition^27–31^. The most commonly downregulated pathways are interferon alpha response and interferon gamma response which are downregulated by palbociclib at 24 hours, trabectedin, ifosfamide, decitabine, capmatinib, olaparib, RMC-4630, and doxorubicin, but upregulated by palbociclib at eight hours and by mirdametinib.

Interestingly, pathway-level effects can differ between our eight- and 24-hour timepoints, such as with interferon gamma response which is upregulated at eight hours with palbociclib but downregulated at 24 hours relative to DMSO control. We also observed pathways uniquely affected by individual drugs. For example, trabectedin affected cholesterol homeostasis and fatty acid metabolism, and the MEK inhibitor mirdametinib was the only drug to affect PI3K-AKT-MTOR signaling (**Table S3**). Together these findings confirm that 3D-MEDS captures pathway-level network adaptation to drug exposure across a diverse compound set, generating the transcriptional response data needed to inform combination prioritization.

### Combination screening reveals heterogeneous drug combination efficacy across MPNST PDX

Next, we evaluated drug combinations using the same 3D-MEDS system, selecting at least one representative from each major therapeutic class included in the single-agent screen, including inhibitors of MEK, SHP2, CDK4/6, HDAC, PARP, and DNA methyltransferase, as well as conventional chemotherapeutic agents. We chose a high dose concentration for each single agent and prepared a range of doses for combination studies (Methods, **Table 1**). We ranked combinations based on mean efficacy of the combination relative to single agents, herein referred to as the ΔAUC (**Fig 4**; Methods). We devised the ΔAUC to summarize the overall impact of a combination compared to each of the single drugs. Though several synergy metrics have previously been established in the literature (e.g., MuSyC,^32^, Bliss^33^, Combination Index or CI^34^), we found that these other metrics were less suitable for EXACT because MuSyC and Bliss were explicitly built to measure drug synergy based on a full matrix of dose response combinations. However, we did calculate these scores and found that they were all significantly correlated with our ΔAUC metric (**Fig. S3 A-C**).

**Figure 4.**
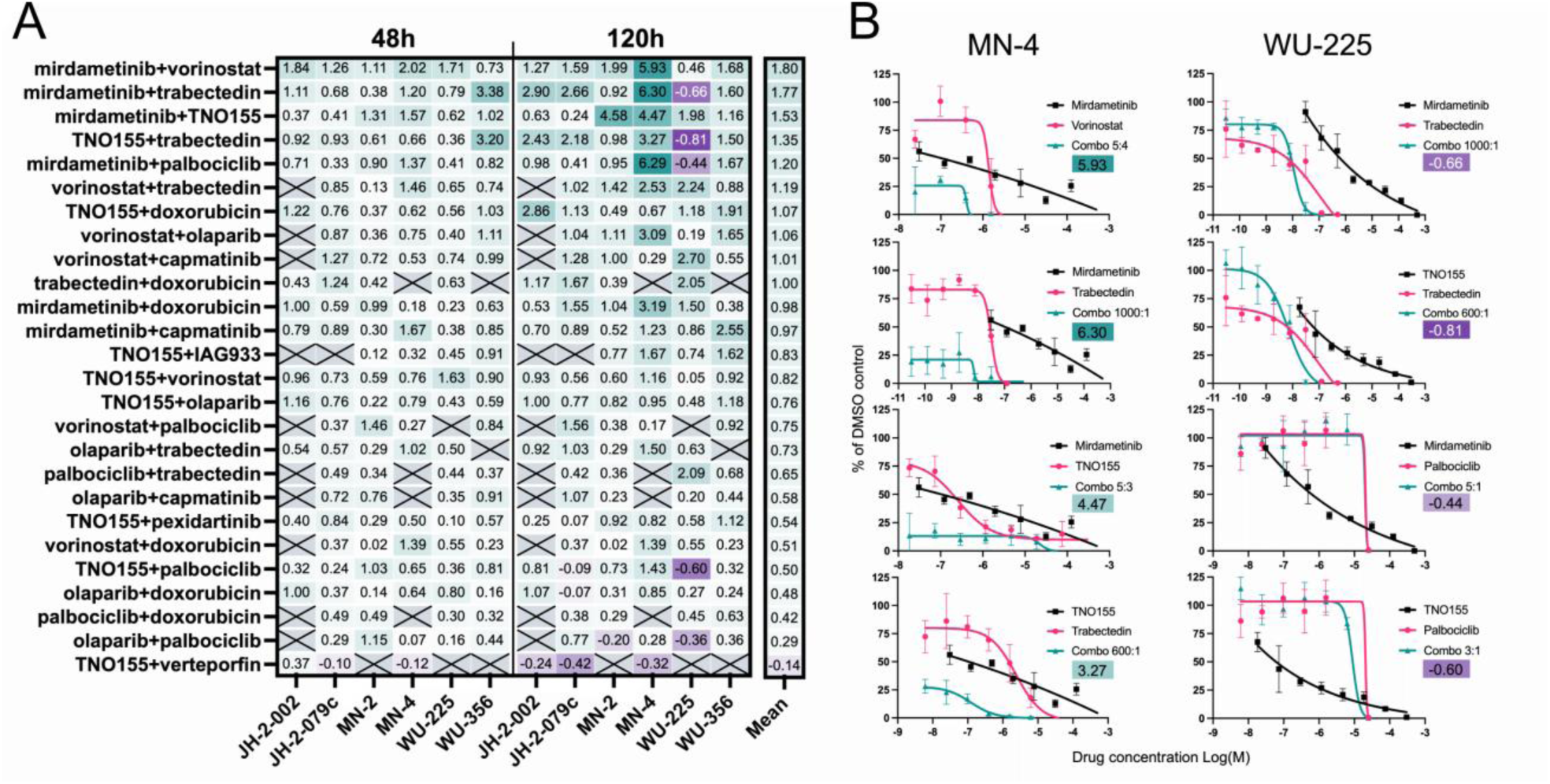
Combination screening of prioritized drug set identifies multiple effective combinations. **(A)** ΔAUC of 25 drug combinations tested in six *ex vivo* MPNST PDX 3D-MEDS for 48 and 120 hours. **(B)** Example dose response curves at 120 hours of MN-4 (left) and WU-225 (right) of some of the highest and lowest ΔAUC values. Combination doses are listed as molar ratios of the first drug to the second drug (e.g., combo 5:4 for mirdametinib+vorinostat means 5 micromoles mirdametinib were administered for every 4 micromoles of vorinostat

Considering we only tested one ratio of concentrations for each drug pair, we found ΔAUC to be more appropriate for this study, despite the fact that it ranked the drug combinations differently than the MuSyC and Bliss metrics (**Fig. S3 D-E.**). Based on our ΔAUC metric, mirdametinib with vorinostat was the most effective combination on average across cell lines tested (**Fig. 4A**). The combination achieved a mean ΔAUC of 1.80 across PDXs (p < 0.001 vs. single agents), with the largest enhanced effect observed in MN-4 at five days. However, it is important to note that the efficacy of these combinations depends on time and the biological context (i.e., the specific MPNST). For example, the combination of mirdametinib with vorinostat is more effective in WU-356 after five days than two days, whereas it is less effective in WU-225 after five days than two days. Overall, MN-4 was the most sensitive to drug combinations (e.g., had the highest ΔAUC values) and WU-225 the least (e.g., had the lowest ΔAUC values) (**Fig. 4B**). These results illustrate the importance of evaluating drug combinations in multiple biological samples even within the same cancer type and at multiple timepoints (**Fig. 4B**).

### Effective drug combinations exhibit overlap upregulated and drug-correlated pathways

We then ranked the combinations by the ΔAUC metric across the three samples for which we collected gene expression perturbation data (MN-2, JH-2-002, and WU-225) and focused on five drug combinations that exhibited a ΔAUC greater than 1, indicating enhanced efficacy relative to the corresponding single agents, depicted in **Table 2**. We used the gene expression statistics calculated in **Fig. 3** to identify additional support of these effective combinations and enable us to prioritize them for additional testing (**Fig. S4**). Specifically, we asked if any pathways associated with sensitivity to one drug (negative correlation in **Fig. 3A**) were upregulated by a second drug (positive enrichment score in **Fig. 3D**). Individual drug results from the top two combinations are shown in **Fig. 5**. Our top-scoring combination includes mirdametinib, which we found to upregulate the inflammatory response pathway across MPNST (**Fig. 5A**). This pathway was also found to be negatively correlated with SHP2i sensitivity across MPNST at 48 hours (**Fig 5B**). The efficacy of this combination was shown in our prior work and suggests that this approach could be used for additional drug combinations ^35^. However, MEK inhibitor plus SHP2 inhibitor combinations would likely result in overlapping toxicities *and* therefore poor tolerability in humans (NCT03989115) ^36^. Our second most effective combination among the three samples we measured via gene expression was mirdametinib with vorinostat. This combination has not been evaluated in previous studies, yet we showed two pathways that served as potential forms of cross talk between the two drugs. **Fig. 5C** shows upregulation of the notch signaling pathway upon vorinostat treatment, and its negative correlation with mirdametinib response at 48 hours in **Fig. 5D**. We observed a similar relationship through WNT–β-catenin signaling (**Fig. S5**). Vorinostat treatment increased the expression of multiple genes in the WNT–β-catenin pathway across the three profiled MPNST models (**Fig. S5A**). Higher baseline WNT–β-catenin pathway activity was also negatively correlated with response to mirdametinib at both 48 and 120 hours (**Fig. S5B**). Thus, both NOTCH and WNT–β-catenin signaling were induced by vorinostat and associated with increased sensitivity to mirdametinib, supporting potential pathway-level crosstalk between the two drugs and providing additional rationale for their combination.

**Figure 5.**
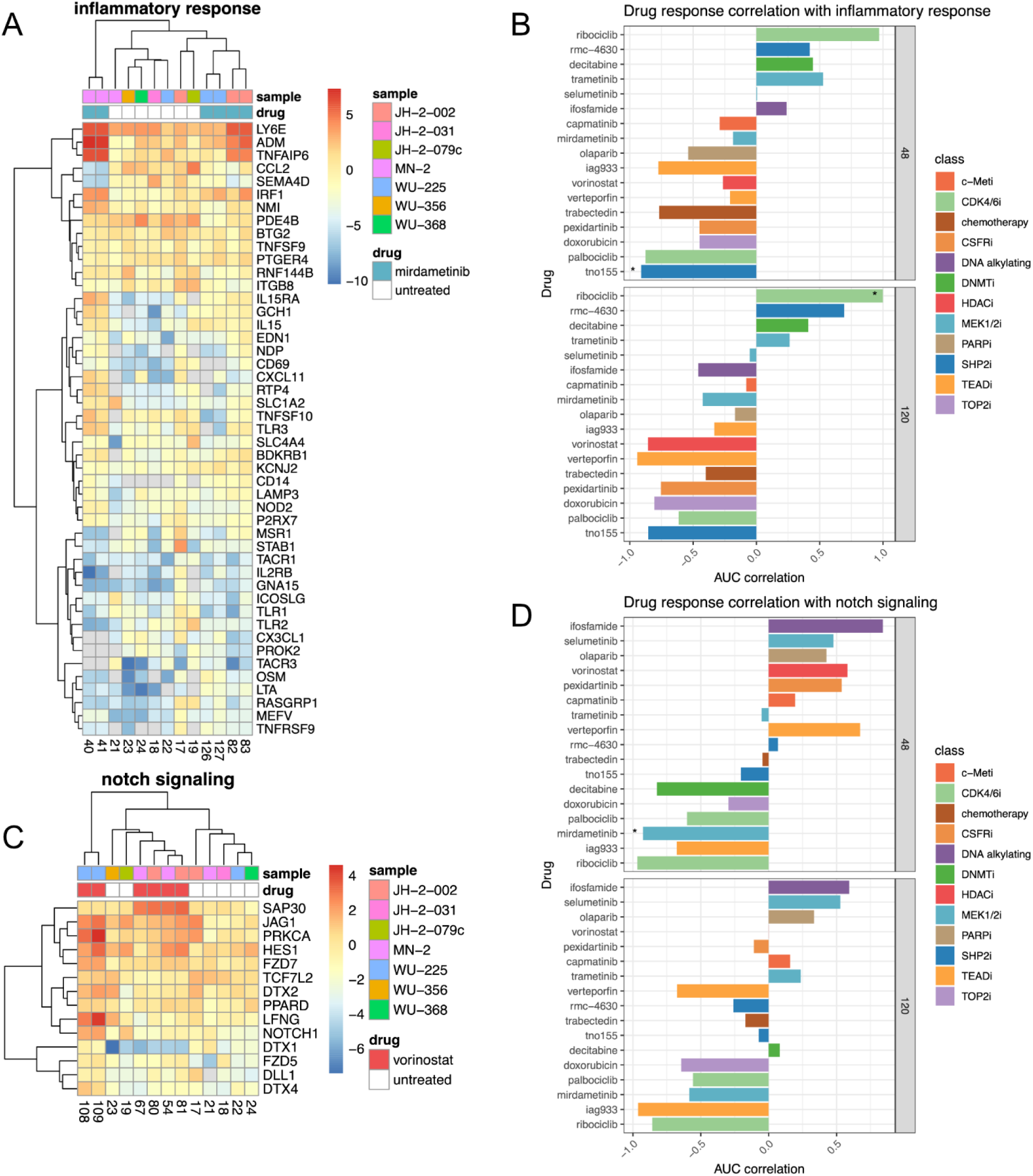
Pathway expression patterns of top-scoring drug combinations. **(A)** Genes in the inflammatory response pathway across untreated MPNST and those treated with mirdametinib. **(B)** Correlation of inflammatory pathway score across untreated samples with the AUC of all drugs at 48 hours (top) and 120 hours (bottom). * Indicates p<0.05. **(C)** Expression of NOTCH signaling genes in untreated MPNST and MPNST treated with vorinostat. **(D)** Correlation of NOTCH signaling scores with drug response across MPNST samples. * Indicates p<0.05.

**Table 2.** Mean deltaAUC of combinations across samples for which we have gene expression data.

| Drug 1 | Drug 2 | mean dAUC |
| --- | --- | --- |
| mirdametinib | TNO155 | 1.58 |
| mirdametinib | vorinostat | 1.40 |
| vorinostat | capmatinib | 1.29 |
| TNO155 | doxorubicin | 1.11 |
| vorinostat | trabectedin | 1.11 |

Since the combination of mirdametinib with vorinostat is not previously reported in MPNST, it warrants further evaluation. We sought to identify further evidence of pathway crosstalk between these two drugs by expanding our correlation analysis to the Cancer Cell Line Encyclopedia and corresponding PRISM drug response dataset, labeling each pathway in the Cancer Hallmarks list by its correlation with drug AUC (see Methods)^3,5^. This approach improves upon our own correlation statistics as we were able to achieve far greater levels of statistical significance across 327 pathways compared to the six for which we measured single dose response data. The results, depicted in **Fig. 6A**, show 11 pathways that are negatively correlated with AUC of a MEK inhibitor with high statistical significance (q<0.05). Specifically, mirdametinib (PD-0325901), MEK inhibitor, is negatively correlated to 7 statistically significant pathways. We then queried these pathways in the MPNST differential expression from our 3D-MEDS system from **Fig. 3D**. The results, shown in **Fig. 6B**, show that many drugs upregulated pathways correlated with MEK inhibitor sensitivity, with vorinostat upregulating the largest number of MEK inhibitor correlated pathways. This relationship adds further evidence to support the combination of a MEK inhibitor (i.e., mirdametinib) and vorinostat *in vivo*.

**Figure 6.**
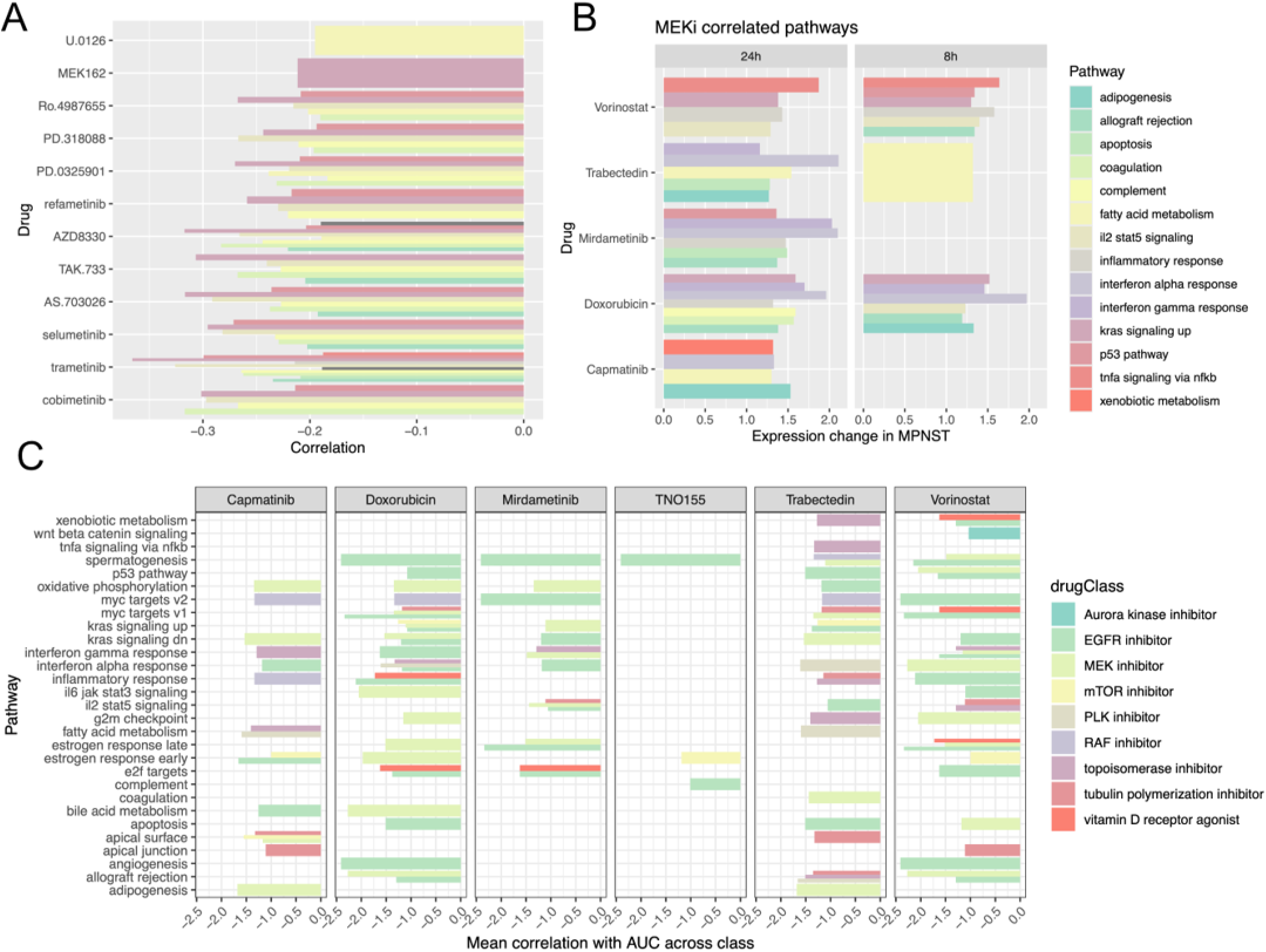
Pathway correlation scores in 2D cancer cell line data provide complementary evidence of drug combination efficacy. **(A)** Cancer Hallmark pathways negatively correlated (q<0.05) with AUC of MEK inhibitors in the PRISM database. X-axis depicts correlation with AUC, y-axis depicts drug name. **(B)** Enrichment of pathways that were negatively correlated with MEK inhibitors up regulated in our own MPNST dataset. **(C)** Drug classes that are enriched in negative correlation scores for a particular pathway of interest. Y-axis selects pathways that are up regulated in MPNST (names across top). X-axis depicts z-score of mean correlation across drugs in that class with the pathway.

Although the PRISM dataset did not include every drug evaluated in our MPNST screen, it contained response data for a much broader range of compounds than the drug screen introduced in this study, enabling us to identify additional drug classes whose activity was inversely associated with the pathways upregulated in our MPNST models. For each pathway identified as up regulated in our MPNST in **Table 2**, we asked if any drug classes were over-represented among the negatively correlated genes using a Kolmogorov Smirnov test (see Methods). The results, depicted in **Fig. 6C**, show additional potential drug classes we can evaluate in MPNST. For example, tubulin polymerization inhibitors and topoisomerase inhibitors were found to be anti-correlated with the MYC targets V1 pathway, which is upregulated by doxorubicin and trabectedin. Additionally, EGFR inhibitors were often anti-correlated with the KRAS signaling down pathway, which was upregulated in our MPNST by mirdametinib, vorinostat, doxorubicin, and TNO155. EGFR inhibition appeared promising in other pre-clinical models due to over-expression of EGFR but a clinical trial of the first generation EGFR inhibitor, erlotinib, in patients with MPNST was not successful ^37^. Based on these observations, we tested one PDX (JH-2-079c) *in vitro* with two different EGFR inhibitors, erlotinib and afatinib. Erlotinib had a modest reduction in cell viability *in vitro* while afatinib was far more efficacious **(Fig. 7A)**. We then treated JH-2-079c *in vitro* with mirdametinib and afatinib in combination **(Fig. 7B)**. This combination showed an enhanced effect compared to either single agent alone. Finally, we tested this combination *in vivo* in the same PDX and the combination slowed tumor growth more than either single agent alone **(Fig. 7C)**. These findings indicate that we can use our transcriptomics data to predict potential secondary drugs from a sample treated with a single agent.

**Figure 7.**
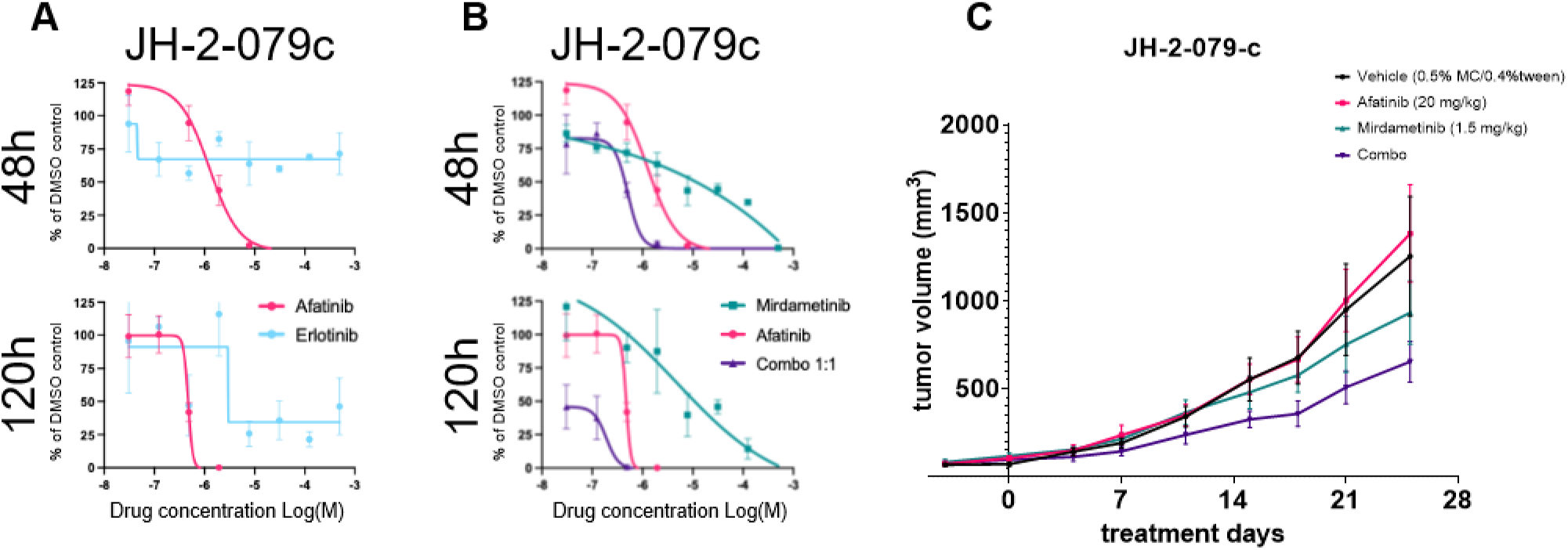
Combination of mirdametinib and afatinib is more effective than single agents alone. **(A)** Dose-response curves show afatinib is more effective in JH-2-079c ex vivo than erlotinib at 48 and 120 hours. **(B**) Dose-response curves for mirdametinib and afatinib either alone or in combination. **(C)** Mean tumor size after 25 days of treatment with mirdametinib, afatinib, or combination in JH-2-079c PDX.

### MEK and HDAC inhibitor combination shows enhanced efficacy in a subset of MPNST PDX

Since our drug sensitivity measurements and pathway statistics indicated that HDAC inhibitors may be effective in combination with mirdametinib in MPNST, we evaluated this combination both *in vitro* and *in vivo* (**Fig. 8A-B**, **Table S1**). Across all *in vitro* samples, the combination reduced viability more than either single agent overall (p=7.72E-15), with effect sizes that were PDX- and time-dependent. The greatest reductions were observed in MN-4 at five days and JH-2-002 at two days, while several other PDX and timepoint combinations did not reach significance. *In vivo*, the combination outperformed single agents in WU-225 (p=6.49E-3), where neither mirdametinib nor vorinostat alone significantly reduced tumor volume. In MN-2, the combination did not outperform single agents (p=2.81E-1); mirdametinib alone was sufficient to significantly reduce tumor volume (p=4.77E-2), potentially masking any additional benefit of the combination at the doses tested. Together, these results demonstrate that EXACT successfully nominated a combination with meaningful *in vivo* activity across both models tested, while the PDX-dependent variability in response both *in vitro* and *in vivo* underscores the importance of evaluating combinations across multiple patient-derived models.

**Figure 8.**
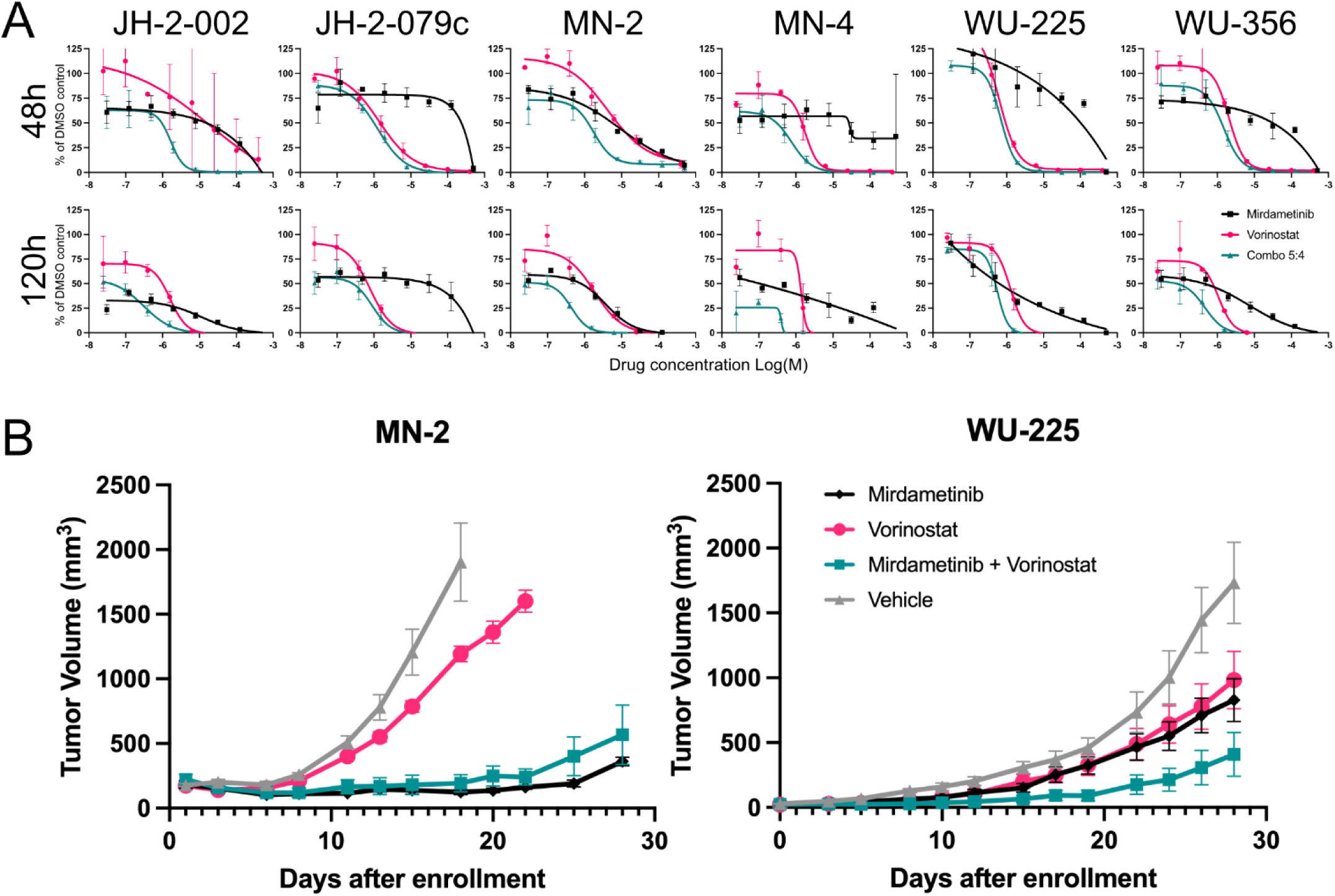
Combination of mirdametinib and vorinostat can be more effective than single agents in MPNST. **(A)** Dose-response curves for mirdametinib and vorinostat either alone or in combination tested in six *ex vivo* MPNST PDX 3D-MEDS for 48 and 120 hours. **(B)** Mean tumor size after 28 days of treatment with mirdametinib, vorinostat, or combination in two MPNST PDX.

## Discussion

In this study, we developed EXACT, an integrated experimental–computational framework for nominating effective drug combinations in rare cancers and applied its utility in the rare and aggressive sarcoma MPNST. The 3D-MEDS platform bridges the gap between low-fidelity 2D assays and low-throughput organoid or *in vivo* models, combining the scalability of standard 96-well screening with the biological advantages of a 3D ECM-rich environment, without requiring specialized fabrication or imaging workflows. By pairing drug sensitivity measurements with baseline and drug-induced gene expression profiling across six PDX lines, EXACT simultaneously captures single-agent activity, network adaptation to drug exposure, and pathway-level evidence for combination prioritization. The resulting datasets, comprising responses (**Table S1**) to 18 single agents and 26 combinations alongside matched transcriptomic profiles (**Table S2**), are shared on the NF Data Portal and provide a foundation for future integration with additional MPNST and rare tumor data.

Among the single agents tested, vorinostat and mirdametinib emerged as the most broadly effective across PDX lines (**Fig. 2A**), consistent with the known roles of HDAC and MEK signaling in MPNST pathobiology and with prior preclinical work supporting both as therapeutic targets in NF1-associated tumors^38,39^. The strong single-agent activity of these compounds across molecularly diverse PDX lines suggests they engage core dependencies in MPNST rather than context-specific vulnerabilities, making them logical anchors for combination strategies. The metabolically active product of irinotecan, SN-38, which is a topoisomerase 1 inhibitor, was also highly effective at reducing viability but since it was only tested in two samples, it is unknown if the effect is more universal. Several combinations involving mirdametinib were among the most effective by ΔAUC (**Fig. 4A**), including pairings with TNO155 and palbociclib, each supported by pathway crosstalk evidence. The MEK inhibitor plus CDK4/6 inhibitor combination is consistent with recent reports of efficacy in patient-derived orthotopic xenograft models, and we reported the MEK inhibitor plus SHP2 inhibitor combination in NF1-MPNST models including those with acquired MEK resistance, lending support to the pathway-based prioritization approach used here ^35,40^. Our CCLE/PRISM analysis additionally nominated EGFR inhibitors as potentially effective partners for several of the drugs tested, a finding with biological plausibility given that EGFR is overexpressed in approximately half of MPNSTs and has been shown to promote tumor formation through STAT3 phosphorylation ^41^. While a clinical trial of erlotinib in MPNST did not demonstrate clinical benefit, the specific combination contexts identified here may represent a more targeted hypothesis worth revisiting in future studies^42^.

The decision to advance mirdametinib and vorinostat to *in vivo* testing was supported by both experimental efficacy data and pathway crosstalk evidence, and prior preclinical work suggesting potential for MEK and HDAC co-targeting in MPNST ^43^. *In vivo*, the combination outperformed single agents in WU-225, where neither agent alone was sufficient to reduce tumor volume, demonstrating that the combination can achieve meaningful activity in contexts where monotherapy fails. The lack of combination benefit over mirdametinib alone in MN-2 may reflect a ceiling effect at the doses tested, as mirdametinib alone was sufficient to significantly reduce tumor volume in that model, though we cannot rule out that the genomic differences between these models also contribute. That said, the genomic differences between these models (e.g., WU-225 carries chr8q gain and p53 loss while MN-2 is p53 wildtype) may also contribute to their differential responses, and a larger panel of PDX would be needed to disentangle dose effects from genotype-specific sensitivity^20^. Nonetheless, the combination showed sufficient *in vivo* activity to support an actively recruiting clinical trial (NCT06693284), representing a direct translational potential of the EXACT framework.

Several limitations of the current implementation should be acknowledged. Drug response and transcriptomic measurements were taken at fixed time points, which may not fully capture the dynamic changes that occur with longer drug exposures, and only one concentration ratio was tested per combination, precluding formal synergy analysis. Not all PDX models were available for every assay, reducing dataset overlap and limiting our ability to fully characterize inter-tumor heterogeneity. The platform currently focuses exclusively on malignant cells and does not capture the influence of stromal or immune components known to modulate drug response in MPNST and other solid tumors, an important direction for future development, including ongoing co-culture studies in our group. Finally, the transcriptomic data used for pathway analysis could be complemented by proteomics, which would provide more direct evidence of shared protein-level mechanisms underlying combination efficacy. Each of these limitations points to concrete opportunities for expanding the platform: broader PDX coverage, multiple treatment timepoints, additional cell types, and deeper multi-omics profiling will strengthen both the experimental and biological rationale arms of EXACT in future studies.

Nonetheless, the EXACT framework represents a significant advance for rare cancer research, providing a physiologically relevant and scalable system to nominate drug combinations with translational potential. By integrating PDX models with a controlled 3D culture environment and leveraging the ongoing expansion of our multi-institutional PDX collection, this platform is well positioned to evaluate a broader range of patient-derived tumors in the future^20^. In addition to established PDX models, incorporating freshly resected and purified primary tumor cells from the operating room may further enhance the clinical relevance of this approach. The paired datasets generated here, drug responses linked to transcriptomic changes and drug combination predictions, will serve as a foundation for future investigations and modeling efforts. This resource will continue to expand, offering the community new opportunities to explore therapeutic strategies and refine methods for predicting effective drug combinations in rare and aggressive cancers.

## Methods

### Drugs

All compounds used in this study are listed in **Table 1** with source and dose information and resuspended in DMSO or H_2_O. Maximum plasma concentration in humans (C_max_) for compounds was determined from literature searches and was converted to molarity. For transcriptomic experiments, we chose a concentration for each compound that was near its C_max_ concentration, if known. For single agent viability studies, we chose a high concentration and performed a four-fold serial dilution for a total of eight concentrations per compound in order to observe responses to a wide range of doses. For combination viability studies, the two agents of a combination were mixed prior to the eight-point serial dilution. Both single agents were tested in parallel with their combination. DMSO or H_2_O was added to control wells to ensure vehicle concentrations were consistent for all experiments.

### 3D-MEDS Establishment and Cell Viability

Patient-derived xenografts (PDX) were established and dissociated as previously described^20^. Single-cell suspensions of dissociated PDX were combined with basement membrane matrix (Matrigel®, Corning, 354234) at a 1:3 (v/v) ratio. 7.5 × 10^4^ cells/well with basement membrane matrix dispensed into white 96-well plates (Corning #3912). Gels were levelled by gentle tapping, incubated for 10 min at room temperature in a biosafety cabinet, then polymerized for 10 min at 37 °C in 5 % CO₂ incubator. To each gel well, 60 µL of pre-warmed custom media (86.5% DMEM, 10% bovine serum, 1.5% HEPES buffer, 1% ITS+ premix, 1% penicillin– streptomycin, 0.001% dexamethasone, and 0.001% glucagon) was added and incubated for 24 h before test compounds were added. For each viability experiment, 1 µL of single agent or pre-mixed combination was added to a well in triplicate. DMSO was also added to vehicle control wells in triplicate with the same final concentration as compound wells. Viability was measured 48 h and 120 h post-treatment with the CellTiter-Glo® 3D assay (Promega, G9681) at a 1:1 reagent-to-well volume, and luminescence was recorded on a microplate spectrophotometer (SpectraMax i3x). Relative luminescence values of experimental wells were normalized to DMSO values after no-vehicle control values were subtracted. Data were analyzed in Prism (GraphPad) with dose response curves generated using nonlinear regression log(inhibitor) vs. response - variable slope and each data point representing mean +/- SD.

### Collagen 3D-MEDS Establishment, RNA Extraction and RNA-Seq Data Processing

Collagen 3D-MEDS were fabricated using high-concentration rat tail collagen I (Corning) and buffered with 10 × DPBS (Fisher, BP399), neutralized to pH 7.4, supplemented with 10% Matrigel (Corning, 356237), diluted to 6 mg/mL concentration, and mixed with 6 × 10^6^ cells/ml. Gels were incubated for 10 minutes at room temperature in a biosafety cabinet, then polymerized for 20 min at 37 °C in 5 % CO₂ incubator. RNA was extracted at 8 h and 24 h post treatment using the TRIzol-RNeasy workflow (Qiagen, 74204) according to the manufacturers’ protocol. Briefly, gels were washed three times in ice-cold PBS, lysed in 1 mL TRIzol, phase-separated with 200 µL chloroform, and the upper aqueous layer (∼450 µL) was mixed 1:1 with 70 % ethanol before loading onto RNeasy columns. After the standard RPE/80% ethanol washes and a 5-min dry spin, RNA was eluted in RNase-free water and quantified by NanoDrop. Total RNA with RIN ≥ 7 was used for library construction using the Illumina TruSeq Stranded mRNA Library Prep Kit with RiboErase (Illumina, San Diego, CA, USA). Libraries were sequenced to a depth of approximately 30 million paired end reads per sample. Raw sequencing reads were processed with the NF-core/rnaseq pipeline (v3.14.0)^44^, implemented with Nextflow. Adapter and quality trimming were performed using Trim Galore, and initial read quality was assessed using FastQC, with results summarized in MultiQC.^44^ Reads were aligned to the human reference genome (GRCh38/hg38) using STAR, and transcript abundance was estimated with Salmon using the GENCODE v40 annotation in alignment-based mode. Raw counts and TPM values were obtained from the quant.genes.sf files, which contain aggregated gene-level expression estimates for downstream analyses. The final outputs included raw and normalized gene counts, transcript abundances, and a comprehensive quality control report.

### Differential expression analysis

DESeq2 was used to calculate differential expression between counts of drug-treated and DMSO control-treated samples ^45^. Genes with adjusted p-values < 0.05 were considered significantly differentially expressed.

### Gene set enrichment analysis

Pathway-level effects of drug treatments were assessed using gene set enrichment analysis (GSEA) of the Hallmark gene set collection (50 pathways) available through the msigdbr R package ^25,46^. Specifically, single-sample GSEA was run first using the gene expression counts from untreated PDX for **Figure 3A**. Pathway scores of untreated samples were correlated with AUC of the each drug in **Figure 2** using Pearson correlation analysis. We then ran the same analysis on the log2(fold-change) in gene expression between drug-treated and DMSO control-treated samples for **Figure 3C**. Pathways with enrichment p-values < 0.05 and FDR q-values < 0.25 were considered significant per the recommendation for GSEA ^46^.

### Dose-response curve fitting

Viability values relative to DMSO controls were divided by 100.00 to put relative viability values on a scale of 0 to 1. Area under the curve (AUC) values were calculated using trapezoidal numerical integration and then normalized by dividing by the absolute dose range. The fit metric R2 was calculated by inputting the difference between relative viability values that were measured and predicted using the Hill-slope equation into the r2_score function from the Python library sklearn.metrics ^47^. Plots were generated using PRISM software.

### Cell line-based pathway correlation

We used the ssGSEA algorithm with the Cancer Hallmarks library to score 327 adherent cancer cell lines in the cancer cell line encyclopedia based ^48^. Next, these similarity scores were correlated with sensitivity area under the curve (AUC) for 1,351 drugs with mechanism of action annotations in the PRISM database ^49^. Finally, enrichment of 83 drug mechanisms of action with at least 6 drugs represented was calculated using this list of 1,351 drugs ranked by Pearson correlation estimates. Drug mechanisms of action with enrichment p-values < 0.05 and FDR q-values < 0.25 were considered significant per the recommendation for DMEA ^50^.

### Drug synergy calculation

To score combinations for potential synergy, we uploaded a CSV file containing percent viability values for each MPNST cell line, timepoint, drug combination, and replicate onto the MuSyC web application^32^. The following parameters were used: Orientation: Emax<E0 and Effect constraint: Unconstrained. Since all dose-response curves reached nearly zero viability by the maximum doses, we focused on the potency metric (i.e., log|alpha|), where a positive log|alpha12| score suggests drug 1 makes drug 2 more potent. Drug combinations were scored based on the average of the log|alpha12| and log|alpha21|. Drug combinations were considered potentially synergistic if both drugs were more potent as a combination rather than as a single agent (i.e., log|alpha12| > 0 and log|alpha21| > 0).

CSV files similar to MuSyC, but which also included drug concentrations, were uploaded to SynergyFinderPlus for Bliss synergy determination ^51^. The following parameters were used: Format: table, Response: % viability and Baseline correction: (complete: correct all values). Bliss scores for each single agent and combination dose were calculated.

Combination Index (CI) values were calculated using CalcuSyn software (Biosoft) as previously described ^34^. ΔAUC values were calculated as the sum of the difference in AUC between single agents and combination treatment divided by the AUC of the combination treatment: [(AUC_1_-AUC_combo_)+(AUC_2_-AUC_combo_)]÷AUC_combo_. This allows us to consider the efficacy of the combination relative to each single agent and scale the resulting score such that positive scores indicate that the combination is more effective on average than the single agents and negative scores indicate that the combination is less effective on average than the single agents.

### Patient-derived xenografts

Patient derived xenografts (PDX) were established by implanting 1x10^6^ cells subcutaneously in the flanks of nude mice. When tumors reached approximately 80-100mm^3^ animals were randomized into a treatment arm or vehicle only control. Mirdametinib was administered at 1.5 mg/kg BID Monday-Friday, vorinostat was administered at 100 mg/kg 3x weekly. Mice were monitored until tumors reached an endpoint size of 2000mm^3^ or 28 days, at which time mice were euthanized.

## Data availability

Data are available upon request on Synapse at https://synapse.org/mpnst. Processing scripts are publicly available on GitHub at https://github.com/PNNL-CompBio/MPNST-PDX-MT/tree/main/EXACT-pipeline.

## Acknowledgements

This work was funded by the American Cancer Society Research Professor Award #123939 to D.A.L.; the CDMRP/ Neurofibromatosis Research Program (NFRP) grant (NF220025, to DAL, DKW, SG, ACH and CAP); and a grant from the Gilbert Family Foundation (to DKW, SG, DAL, ACH and CAP). The authors acknowledge the Sachs lab at University of Minnesota for use of their microplate reader.

## Author information

### Contributions

R.C.S. generated all the *in vitro* single agent and combination data and contributed to manuscript writing. A.T.L. determined *in vitro* drug concentrations and dilutions, analyzed viability data, and contributed to manuscript writing. B.B.G. analyzed RNA-seq and viability data, generated drug sensitivity predictions, and led the writing of the results and discussion sections. Y.L., Z.S., E.R., K.B.W., D.B., K.X., X.Z., Y.M., J.J., L.B.F, and Y.Z. contributed. D.A.L., C.A.P., A.C.H., D.K.W., S.J.C.G. provided supervision, read and edited the manuscript.

## Supplementary information

**Figure S1.**
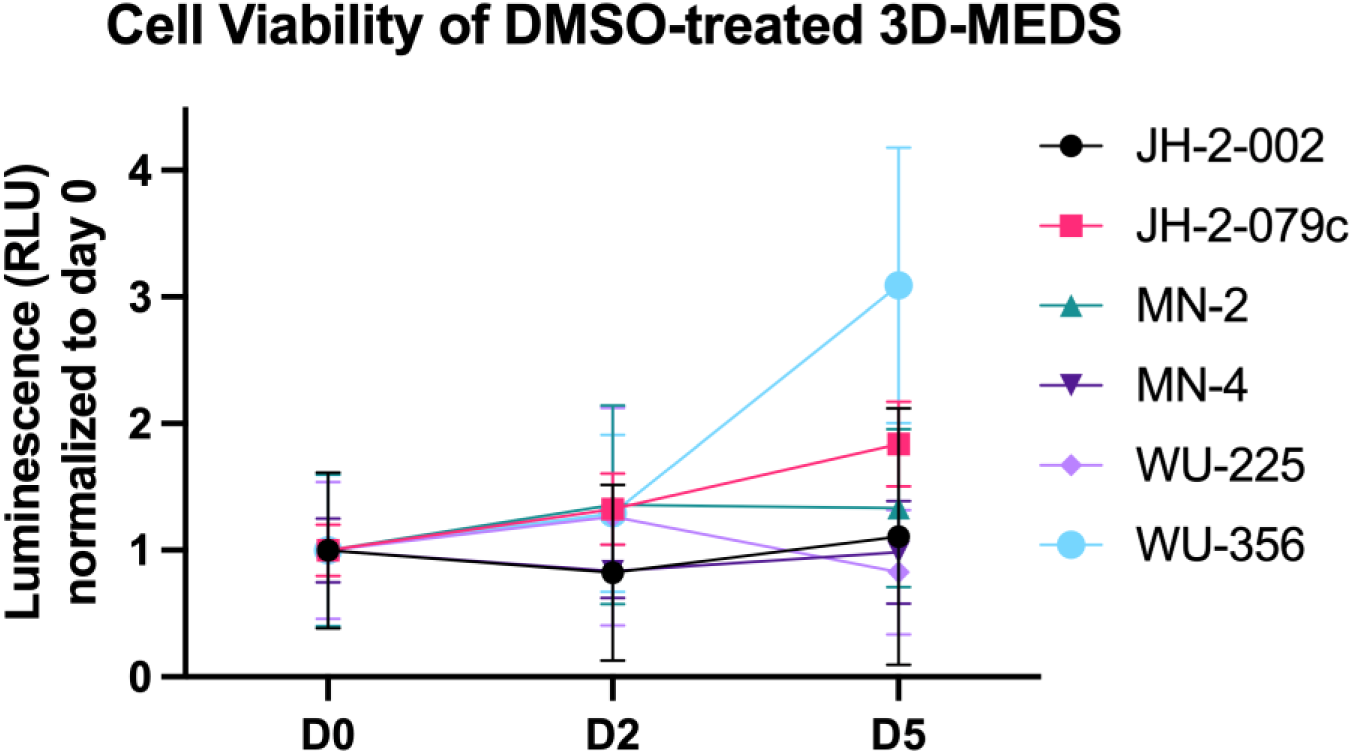
3D-MEDS platform enables cell viability and proliferation. Six MPNST PDX were assembled in the 3D-MEDS platform and treated with DMSO for five days. CellTiter Glo was used to assess cell viability at each timepoint with a luminescent readout. Five of the six PDX had an increase in luminescence on either day 2 or day 5 suggesting proliferation and all PDX were viable on day five. Luminescence values were normalized to mean of day 0 timepoint for each PDX with data points representing mean normalized luminescence values ± SD (n ≥ 6).

**Figure S2.**
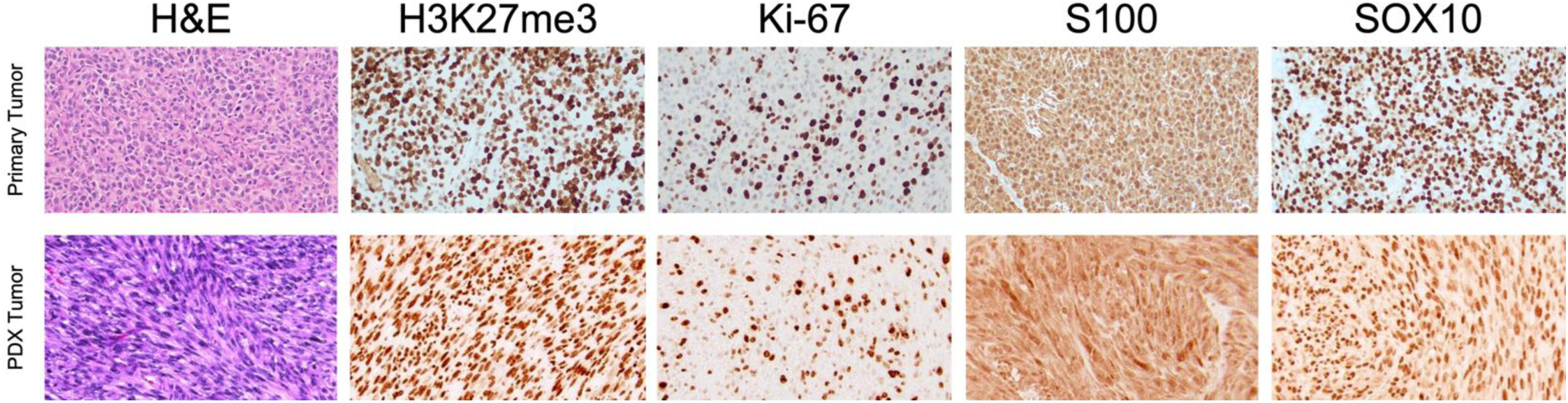
Immunohistochemistry staining of matched MN-4 primary tumor and patient derived xenograft tumor. MN-4 primary patient tumor (top) and patient derived xenograft tumor (bottom) were stained for the following markers: H&E, H3K27me3, Ki-67, S100 and SOX10. Both tumor types show retained expression of H3K27 trimethylation, Ki-67, S100 and SOX10.

**Figure S3.**
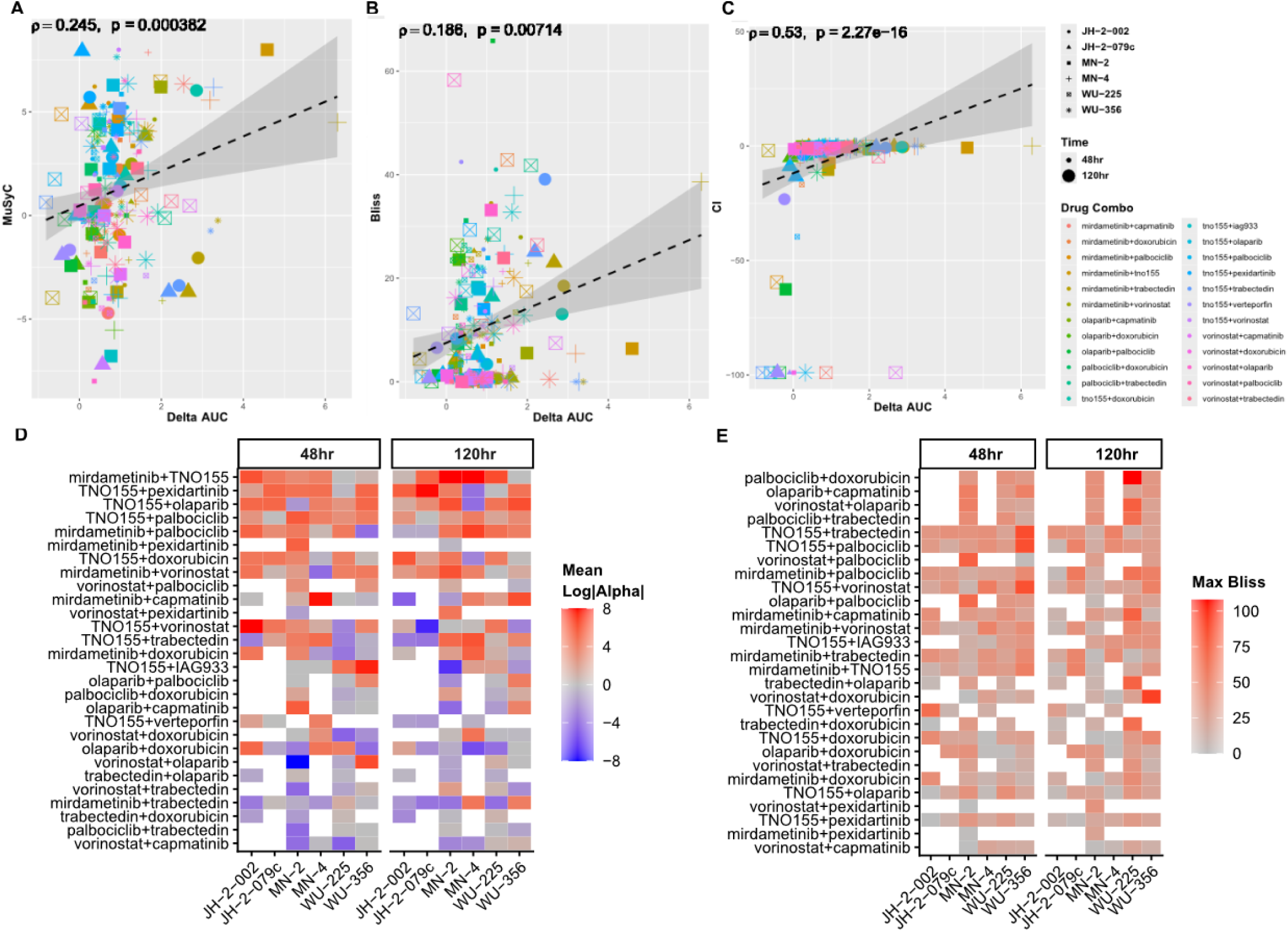
Synergy scores affect drug combination rankings. Correlations between ΔAUC and synergy scores: MuSyC mean log|alpha| (**A**), max Bliss scores for tested drug concentrations (**B**), and negative combination index (**C**). Drug combinations ranked by mean of either MuSyC mean log|alpha| (**D**) or max Bliss scores for tested drug concentrations (**E**).

**Figure S4.**
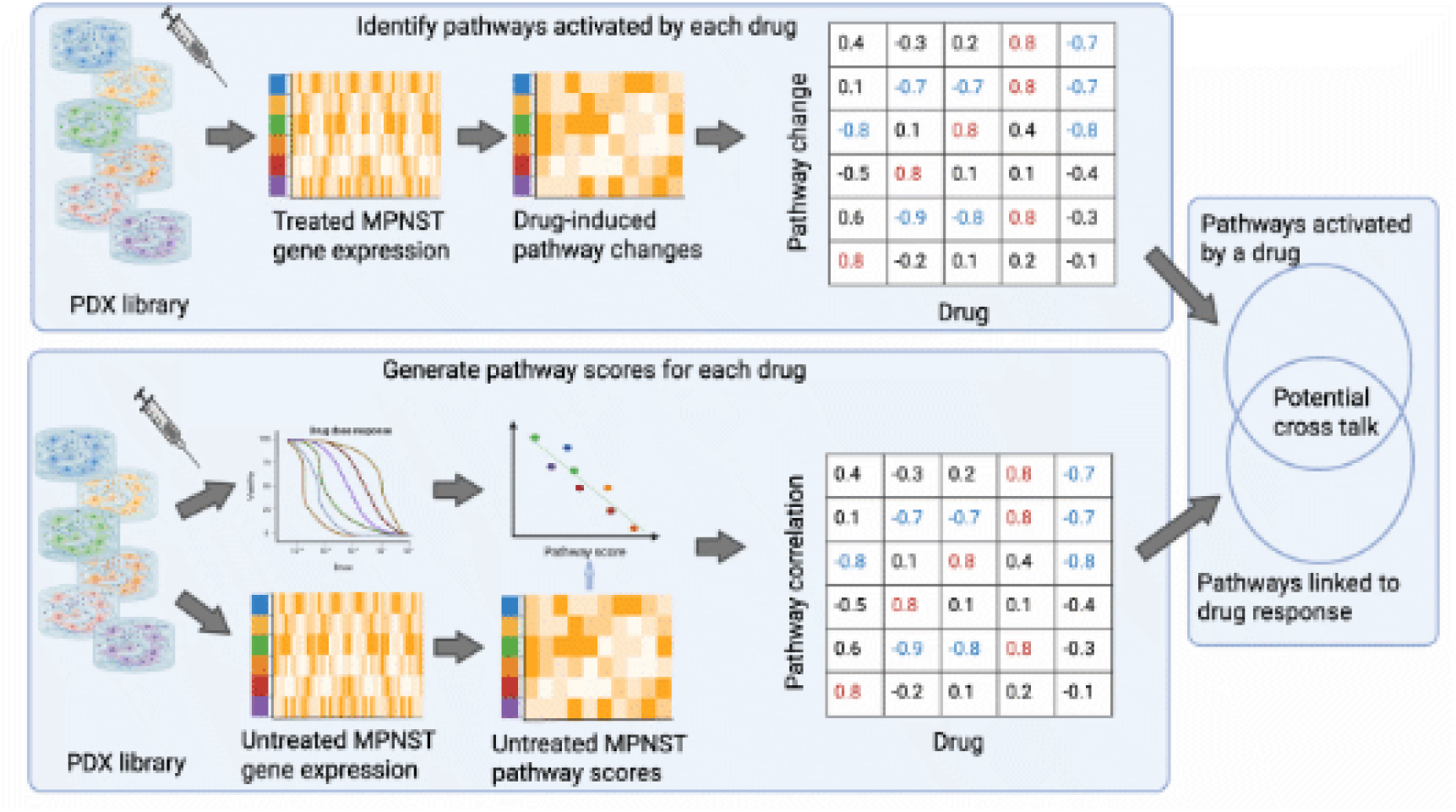
Schematic describing the pathway-based drug analysis. Top: Gene expression changes from single agent treatments are mapped to the Cancer Hallmarks signature database to identify specific signatures that are activated upon drug treatment. Bottom: published gene expression data is reduced to the cancer Hallmark pathways using ssGSEA and compared to single agent AUC values to identify specific pathways correlated with drug sensitivity (indicated by a negative correlation). Right: pathways that are up regulated upon one drug treatment (top) and correlated with drug sensitivity with a second drug (below) indicate potential cross talk and suggest that these two drugs could work together.

**Figure S5.**
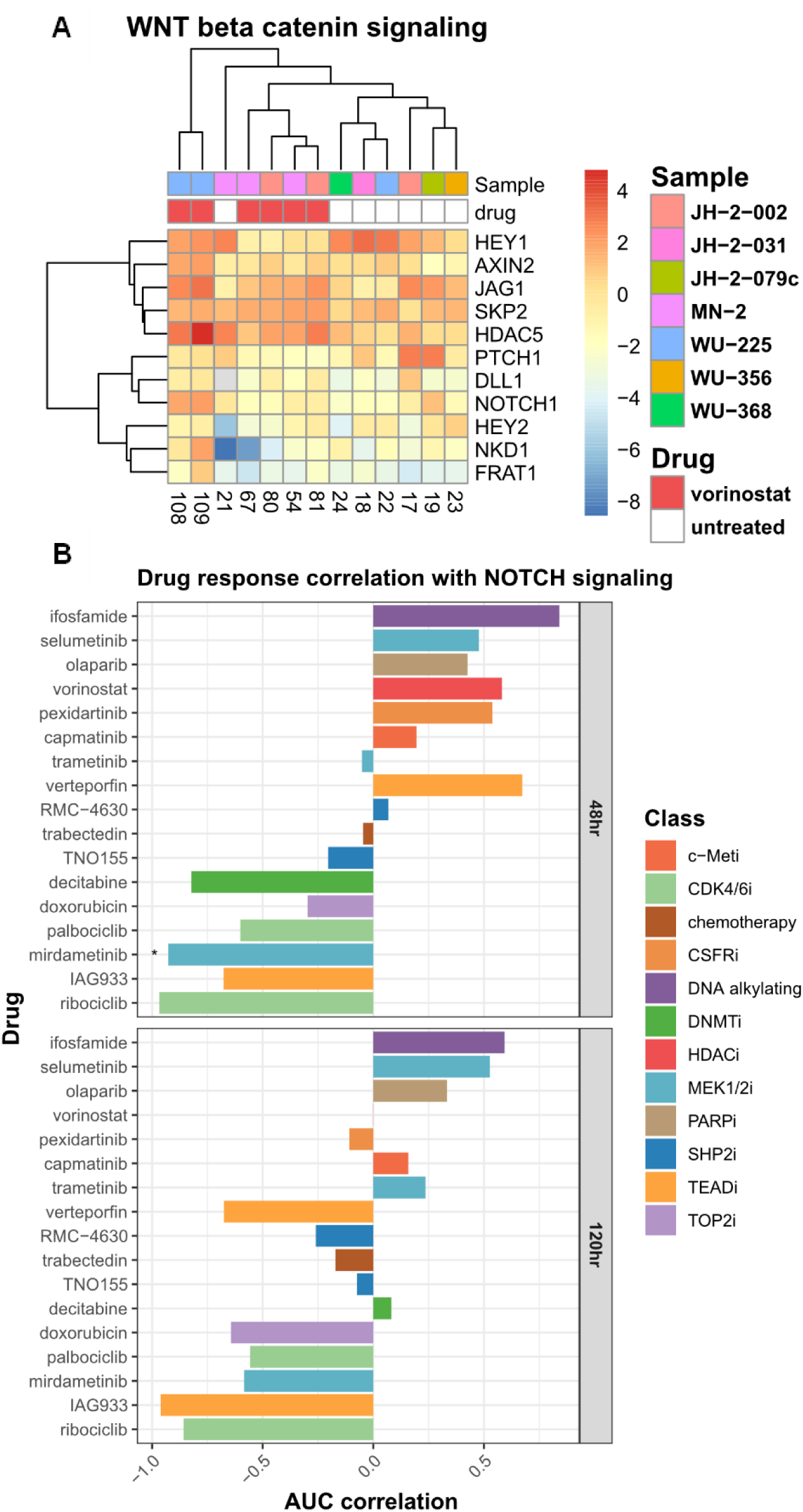
WNT–β-catenin pathway expression and drug-response correlations. (A) Expression of genes in the WNT Beta Catenin Signaling pathway across untreated MPNST samples and samples treated with vorinostat. (B) Correlation of WNT Beta Catenin Signaling pathway scores across untreated samples with the AUC of all drugs at 48 hours (top) and 120 hours (bottom).

**Table S1. Viability**

Viability data normalized to DMSO controls after single or combination drug treatments, area under the curve (AUC) values, ΔAUC values, and PDX tumor sizes.

**Table S2. Differential expression results**

Differential expression between drug-treated samples and DMSO controls.

**Table S3. GSEA results**

Gene set enrichment analysis results using MSigDB gene set collections.

**Table S4. DMEA results**

Drug mechanism enrichment analysis results, including drug sensitivity correlations

